# Drivers of asymmetrical insect invasions between three world regions

**DOI:** 10.1101/2023.01.13.523858

**Authors:** Rylee Isitt, Andrew M. Liebhold, Rebecca M. Turner, Andrea Battisti, Cleo Bertelsmeier, Rachael Blake, Eckehard G. Brockerhoff, Stephen B. Heard, Paal Krokene, Bjørn Økland, Helen Nahrung, Davide Rassati, Alain Roques, Takehiko Yamanaka, Deepa S. Pureswaran

## Abstract

The geographical exchange of non-native insects can be highly asymmetrical, with some world regions ‘exporting’ or ‘importing’ more species than others. Several hypotheses have been proposed to explain such asymmetries, including differences in propagule pressure, environmental features in recipient regions, or biological traits of invaders. We tested aspects of these hypotheses in the context of the exchange of non-native insects between North America, Europe, and Australasia. Europe was the dominant exporter of non-native insect species between the three regions, with most of this asymmetry arising prior to 1950. The European dominance could not be explained by differences in import value, source species pool sizes, or native plant richness in the recipient regions. We identified that the introduction of non-native plants, driven in part by European colonization, best explains the asymmetrical exchange of non-native insects between our focal regions.

## Introduction

Biological invasions can transform ecosystems, with serious ecological, economic, and cultural consequences worldwide. Non-native insects have been implicated in displacing native species, altering the composition of ecological communities, damaging economically important plants including trees and food crops, vectoring diseases, and more^1,2^. Biosecurity measures of various kinds, such as phytosanitation and surveillance programs, are implemented in many countries to minimize new non-native insect establishments; these measures are informed in part by knowledge of the most likely invasion pathways and the taxa that pose the greatest risk^3^. Overall risk associated with the invasion of an individual insect species is determined both by the probability that the species becomes established and the damage that it is likely to cause once established. In turn, a species’ establishment probability is affected both by the likelihood that it is transported to the recipient region and the probability that arriving populations successfully sustain themselves^4^.

An intriguing aspect of invasion risk is that some regions “export” disproportionately more non-native insects during biotic exchange than others. For example, considerably more phytophagous forest insects have invaded North America from Europe than the reverse^5^. The question of why such asymmetries occur has fascinated ecologists for decades, with several mutually compatible hypotheses offered^5–8^:

1. *The source species pool size hypothesis:* the number of invaders from a region should be proportional to the number of species in that region^8^. Using simulations, Seebens et al.^9^ showed that the total number of species that will eventually establish into a region is determined mostly by the size of the pool of potential invaders. However, they also note that the pool of potential invaders is a subset of the total species pools and depends on many interacting factors including species distributions and abundances, environmental suitability, and propagule pressure. Thus, species richness in a donor region is an imperfect proxy for the size of the pool of potential invaders coming from that region.
2. *The propagule pressure hypothesis:* differences in the rate of establishments between regions are driven by differences in arrival rates caused by differences in the magnitude of trade or other invasion vectors^7,8,10^. International trade dominates as a driver of propagule pressure, though it is partly counteracted by biosecurity measures^10^. Propagule pressure may be the most important determinant of establishment success, as high propagule pressure is likely required to overcome environmental and demographic stochasticity, and Allee effects such as mate-finding failure^11^. Recent literature suggests that trade patterns correlate well with the establishment rates of non-native insect species^12–14^. In addition, events such as world wars and accelerating globalization which affected arrival rates have also left their mark on temporal patterns of species establishmens^15,16^. However, as trade increases, the per-shipment probability of transporting new species does not scale linearly. Additional shipments do not bring entirely new assemblages of species, but rather ‘samples’ of species pools that have previously been sampled from. Species that are abundant in a donor region are more likely to be transported to a recipient region and become established earlier than less abundant species, leading to a decline over time in the number of novel species that are introduced to the recipient region. Consequently, effects of increasing trade may be attenuated by the gradual exhaustion of donor species pools and the saturation of invaders in the recipient region^10,12^.
3. *The recipient environment hypothesis:* differences in the environments of recipient regions (e.g., climate, ecology, or existing biota) may promote or inhibit invasion. The lack of suitable climates or host plants may explain many failed establishments of introduced insects. Furthermore, approximately 75% of herbivorous insects exhibit a degree of host-specificity^17^, thus successful establishment may depend on the phylogenetic similarity of potential host plants between the native and non-native habitats^11^. Diversity of both native and non-native host plants appears to be a strong driver of insect invasions^18^. The role of non-native plants in facilitating the establishment of insects is particularly noteworthy given that European colonization promoted the worldwide establishment of European plants and increased the similarity of floral composition between empires and their colonies^19^.
4. *The invader quality hypothesis:* differences in the biological traits of insects native to some regions may make them better at invading or competing than those native to other regions^7,8^. For example, life history traits such as asexual reproduction (e.g., parthenogenesis) and sib-mating are likely to reduce mate-finding failure in incipient populations, strong dispersal abilities may help individuals find suitable host plants, and species that are effective at colonizing disturbed habitats may be particularly effective invaders^11^.

In their influential 1996 paper, Niemelä and Mattson^5^ favored the recipient environment and invader quality hypotheses to explain a relative overabundance of European phytophagous insects in North America compared to the converse. They proposed the existence of a ‘European crucible’ resulting from repeated and extensive glaciations which pushed species into highly isolated refugia. The European crucible may have led to the extinction of many plant genera (particularly of trees), reducing the availability of suitable niches for phytophagous insects. Furthermore, repeated isolation and release from refugia may have made European species superior competitors in disturbed and fragmented habitats, possibly increasing their ability to invade novel habitats^5^. However, aspects of Niemelä and Mattson’s European crucible hypothesis are controversial; when non-native insects from all world regions are taken into consideration, Europe has approximately as many established non-native forest insects as does North America, and does not appear to be particularly resistant to invasion^20^. This suggests that the asymmetry in non-native forest insects exchanged between Europe and North America may be a consequence of the specific invasion pathways and histories between these two regions.

We considered the above hypotheses in the context of asymmetric exchange of all non-native insect taxa between three world regions that engage in significant anthropogenic interactions and exchange of species: North America, Europe, and Australasia (limited to Australia and New Zealand in our analyses), with the goal of testing predictions arising from the source species pool size, propagule pressure, and recipient environment hypotheses:

1. Prediction for the source species pool size hypothesis: donor regions with greater richness of native insect species will export proportionally more insects to recipient regions.
2. Prediction for the propagule pressure hypothesis: differences in trade volume, using inflation-corrected import values as a proxy, will drive differences in the exchange of non-native insects between region pairs among Europe, North America, and Australasia.
3. Prediction for the recipient environment hypothesis: recipient regions with greater richness of native or non-native plants may provide greater niche diversity, and thus facilitate the establishment of non-native insects.

Due to the difficulty of quantifying the ‘invasiveness’ of regional source species pools, we only indirectly address the invader quality hypothesis as a possible explanation for variation that cannot be convincingly attributed to other causes. An additional difficulty is that records of first discovery may be a poor proxy for dates of establishment, due to the time lags that usually occur between establishments and discoveries^21^. We addressed this concern by using Poisson process models that allowed prediction of discovery time lags and the numbers of annual establishments that best fit records of first discoveries. By drawing on recent literature and combining datasets of annual discoveries of non-native insects and annual import values, we have confirmed that disproportionate numbers of European insects have established into North America and Australasia, across all taxa. However, our analyses do not support the notion that differential import values or native plant diversity in the recipient regions are shaping these asymmetries. Instead, we propose that the disproportionate invasion of European insects into North America and Australasia can be explained by European colonization, which facilitated the establishment of European plants and their associated insects and changed floral composition such that European insect invaders were more likely to find suitable host plants.

## Methods

### Datasets and world regions

Insect establishment data were based on the International Non-native Insect Establishment database^22^, supplemented by several other online datasets^9,23–26^. We used an automated taxonomic cleaning script^27^ using the GBIF^26^ API to standardize species names (merge synonyms and correct misspellings).

We worked with a subset of the establishment database that allowed us to compare non-native insect discoveries between donor and recipient regions. The chosen regions were North America (NA), Europe (EU), and Australia and New Zealand combined into an Australasian region (AU). Note that there were minor mismatches in the spatial extents of these regions depending on context. For example, as a donor region, Australasia included Papua New Guinea, but as a recipient region, it only included Australia and New Zealand because we did not have non-native insect discovery records for Papua New Guinea.

For all analyses except the regional counts of non-native insects reported in Table 1, we excluded discovery records where: (1) species had native ranges spanning multiple biogeographic regions (e.g., Holarctic or cosmopolitan species); (2) the native ranges and establishment regions were the same (indicating species spread within these regions); (3) the establishment was limited to “indoors” (e.g., greenhouses); or (4) the establishment was a result of intentional introduction. This left us with a dataset of 2,324 non-native insect discovery records across the six pairwise routes between North America, Europe, and Australasia. For the regional counts of non-native insects reported in Table 1, we only excluded intentional and indoor-only establishments.

**Table 1.** Approximate numbers of described native and non-native vascular plants and insects in North America (NA, north of Mexico), Europe (EU), and Australasia (AU). Counts of non-native insects are from all world regions in our dataset (excluding intentional and indoor-only introductions). All other counts are obtained from literature or taxonomic databases (see citations). We used log-likelihood ratio goodness-of-fit tests (*G*-tests) to assess the equality of counts between regions for each category.

|  | Vascular plants |  | Insects |  |
| --- | --- | --- | --- | --- |
|  | Native | Naturalized non-native (extra-continental)<sup>35</sup> | Native | Non-native (all origins) |
| North America | 15,447<sup>36</sup> | 3,513 in total<br>(1,839 from EU<br>364 from AU) | 87,107<sup>37</sup> | 3,364 |
| Europe | 16,000<sup>38</sup> | 1,883 in total<br>(759 from NA<br>163 from AU) | 94,000<sup>39</sup> | 1,439 |
| Australasia | ~ 21,826*<br>(19,324 in Australia<sup>40</sup>,<br>2,502 in New Zealand<sup>41</sup>) | 3,371 in total<br>(1,322 from EU<br>775 from NA) | 81,667<sup>42,43</sup> | 2,146<br>(1,004 in Australia,<br>1,543 in New Zealand) |
| *G* (*df* = 2) | 1,363 (*p* < 0.001) | 601 (*p* < 0.001) | 955 (*p* < 0.001) | 813 (*p* < 0.001) |

\* Assuming few native species are common to both Australia and New Zealand due to high endemism of plants in New Zealand<sup>30</sup>.

Trade data were obtained from the TradeHist database^28^, modified with modern ISO-3 country codes. The TradeHist database describes the annual value of trade goods from 1827–2014 in British pounds sterling (not corrected for inflation) flowing from origin to destination countries. We corrected all trade values for inflation relative to 2020 based on the annual percent change of the UK retail prices index^29^. We grouped the origin and destination countries into the same regions as above (North America, Europe, and Australasia), with some minor unavoidable differences where national borders did not follow biogeographic boundaries. From these groupings of countries, we created a subset of the TradeHist database representing the six directional pairwise routes between North America, Europe, and Australasia by summing annual trade value across all countries within each region. Records of trade between countries within each of the resulting biogeographic regions were dropped.

### Analyses and Poisson process models

We obtained estimates of species richness for native vascular plants, non-native vascular plants, and native insects in our focal regions from a variety of literature sources (see citations in Table 1). However, we could not find an estimate of native plant richness for the combined geographical area of Australia and New Zealand. Because endemism of native plants in New Zealand is estimated to be greater than 80%^30^ we used the sum of native plant richness in Australia and New Zealand, based on the assumption that relatively few species would be common to both regions. The total numbers of discovered non-native insects in each region were calculated as the sums of unique non-native insect species binomials in each region’s discovery records, after excluding intentional and indoor-only introductions but prior to any further exclusions. We used *G*-tests (log-likelihood ratio goodness-of-fit tests) to test the null hypothesis of equal species richness of plants and insects, both native and non-native, between our three regions.

To investigate simple spatial asymmetries in non-native insect discoveries, we tallied the number of first discoveries of non-native insects for each of the six directional pairwise routes between North America, Europe, and Australasia. We investigated possible explanations based on plant and insect richness by calculating the proportions of non-native insect discoveries to (i) native insect richness in the donor community, (ii) native plant richness in the recipient community, and (iii) the richness of non-native plants which established from the donor to the recipient. To compare the variation between these proportions and non-native insect discoveries, we calculated the coefficients of variation (CV; standard deviation divided by the mean) separately for each metric.

To further investigate temporal patterns of non-native insect establishments between the six routes, accounting for lags between species establishments and discoveries, we used a Poisson process modelling approach modified from Costello et al.^31^ and Morimoto et al.^32^. These models estimated the lag between establishments and discoveries, predicted the annual establishments necessary to fit to observed discoveries given the lag estimates, and (optionally) modelled reductions in the per-unit-import rate of non-native insect establishments due to the saturation of finite species pools. Because the Poisson-process models depend on dated discovery data, we excluded discovery records with unspecified dates. This left us with 1,945 dated records (84% of the dataset).

We omitted an intercept term in our models, forcing them to account for all establishments as a function of imports. We modelled the saturation of establishments as a non-linear (squared) probability of establishment based on the assumption that the invader pool will be rapidly depleted of its more numerous and/or best invaders. These modifications were necessary to produce good fits to our data – initial attempts to use the same models as in Morimoto et al.^32^ resulted in nonsensical parameter estimations and poor fits in most cases. Our full model was as follows:

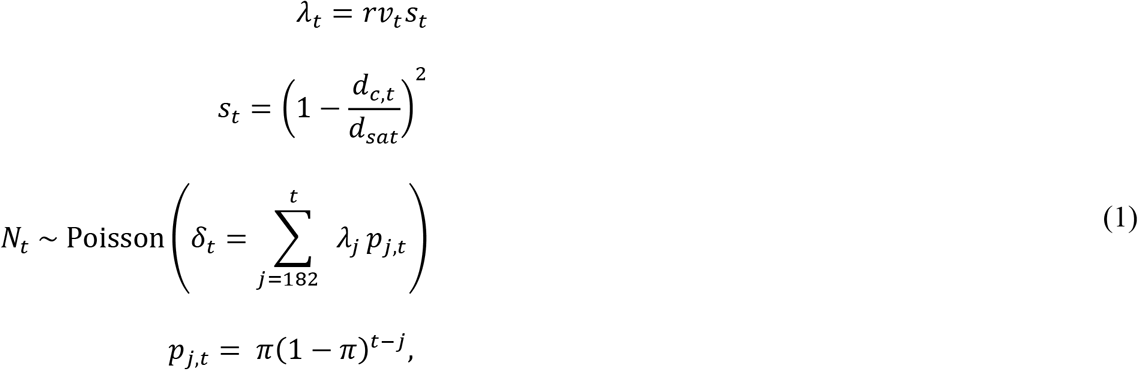

where:

*λ_t_* is the predicted number of new non-native establishments in year *t*,
*r* is the number of species established per billion pounds sterling (prior to saturation),
*v_t_* is the value of imports (2020 billion pounds sterling) in year *t*,
*s_t_* is the per-species probability of establishment in year *t*,
*d_c,t_* is the (observed) cumulative species discovered by year *t*,
*d_sat_* is the number of discoveries after which new establishments cease (saturate),
*N_t_* is the actual number of non-native discoveries in year *t*,
*δ_t_* is the predicted number of non-native discoveries in year *t*,
*p_j,t_* is the probability that a species which established in year *j* will be discovered in year *t*,
and *π* is the annual probability of discovery.

The cumulative sum of discoveries (*d_c,t_*) was calculated by summing the number of annual discoveries from the first year of records (1827) to year *t*, inclusive. We used the sum of discoveries instead of establishments for modelling the saturation of species pools because discovery sums could be easily calculated from the original data. The main drawback to this technique was that it slightly complicated the interpretation of the saturation parameter (d_sat_): rather than being a predicted maximum number of established species, it was the predicted number of discoveries at which point the maximum number of established species had been reached.

All analyses were performed in R 4.1.0^33^. We fit the models to observed annual discoveries (*N_t_*) for each combination of donor and recipient region, minimizing the maximum likelihood as described by Morimoto et al.^32^:

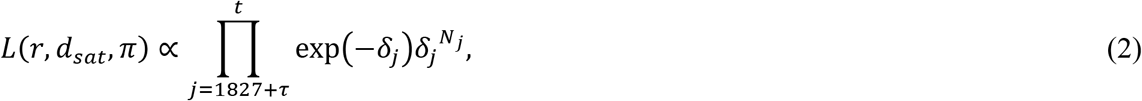

where *τ* =20 as “preservation years” to prevent fitting the model to species that established prior to 1827 (the first year of discovery records in our database) but were discovered after 1827. Without these “preservation years”, *δ_t_* (the predicted number of discoveries in year t) may be underestimated near the start of the dataset because there will be a lack of prior years of predicted establishments from which to model the lagged discoveries^32^. We also used a reduced model which omitted the saturation effect from Eq. (1), effectively making *s_t_* a constant with a value of 1. We then removed the associated parameter (*d_sat_*) from Eq. (2). This “without saturation” model was otherwise identical to the full model.

For parameter estimation, we set lower and (in a few cases) upper bounds on each parameter using the ‘L-BFGS-B’ method^34^. We bounded the rate of establishments (*r*) to ≥ 0.005 non-native species per billion pounds sterling, and the annual probability of discovery (π) to between 0.0125 and 0.95 (corresponding to 1.05 – 80 years of discovery lag). For the saturation term (*d_sat_*), we set the lower bound to the cumulative number of dated discoveries, which varied for each of the six routes (1121 for EU to NA, 205 for NA to EU, 349 for EU to AU, 70 for AU to EU, 74 for NA to AU, and 53 for AU to NA), with no upper bound. We fit both the full and reduced models (the latter lacking the saturation term) to each establishment route and selected the one with the lowest Akaike information criterion (AIC) value.

The *R* function *optim* was employed for all the parameter estimation in Poisson process models (R Core Team, 2022). The confidence intervals are approximately calculated using the inverse of the Hessian matrix evaluated at the last iteration in the optimization process. For parameters with lower or upper bounds, we truncated the confidence intervals to the parameter estimation boundaries.

Observed annual discoveries and model predictions of annual establishments and discoveries were plotted as cumulative curves versus cumulative import value, based on Levine and D’Antonio^12^.

## Results

### Regional differences in cumulative discoveries, plant richness, and species pools

Species counts of vascular plants and insects, both native and non-native, differed significantly between North America, Europe, and Australasia (all *p* < 0.001; Table 1). Counts of non-native plants and insects are particularly variable between regions and follow similar patterns: Europe has approximately half as many non-native plants as North America or Australasia, 57% fewer non-native insects than North America, and 33% fewer non-native insects than Australasia (Table 1).

European insects accounted for 80% of the discoveries of non-native insects exchanged between Europe, North America, and Australasia. The bidirectional exchange of species between Europe and North America was highly asymmetrical, with 6.1 times more European insects discovered in North America than the converse. Nearly the same relationship was seen between Europe and Australasia, with 5.8 times as many European insects discovered in Australasia than the converse (Fig. 1a).

**Figure 1.**
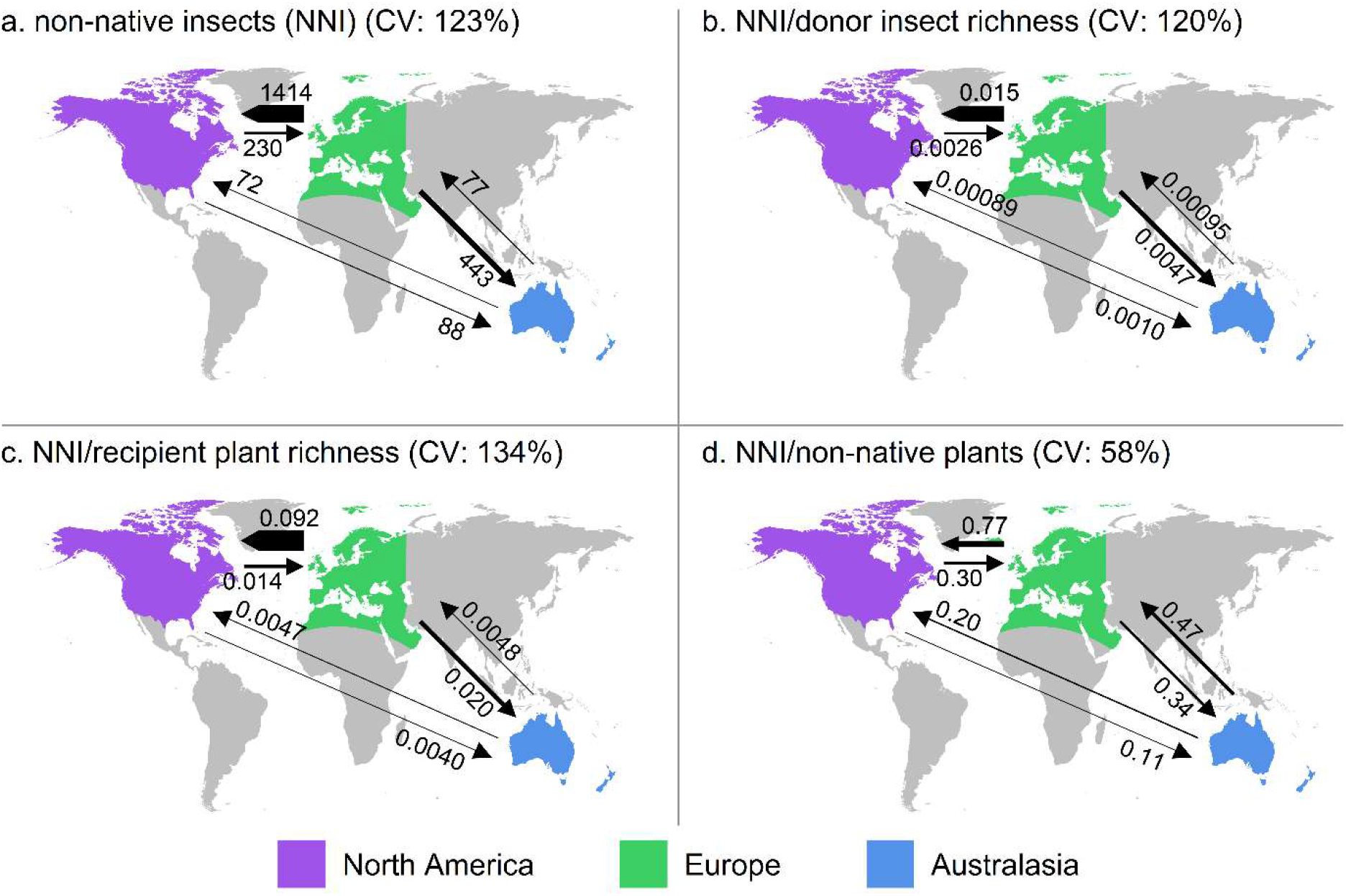
Counts of discovered non-native insects exchanged between North America, Europe, and Australasia (a) and the proportions of these counts to the richness of native insects in the donor regions (b), the richness of native plants in the recipient regions (c), and the number of non-native plants established from the donor to the recipient regions (d). Proportions were calculated from the values in Fig. 1a and Table 1. Arrow widths are proportional to the counts/proportions. CVs are the coefficients of variation (standard deviation divided by the mean) for each of the six counts/proportions per panel.

The relative magnitudes and coefficients of variation for non-native insect discoveries were similar regardless of whether they were based on simple counts of non-native insects, their proportions to donor species pool sizes, or their proportions to recipient native plant richness. In contrast, expressing non-native insect discoveries as proportions of the numbers of non-native plants established over the same routes reduced the apparent asymmetries between donor and recipient pairs, and consequently reduced the coefficient of variation (Fig. 1).

### Spatiotemporal analyses and Poisson process models

Per-unit-import establishments of European insects into North America began to slow in the 1950s, indicated by highly convex plots of cumulative establishments/discoveries versus cumulative imports. The best model included a saturation effect. By 1950, approximately 75% of non-native insects from Europe had already established into North America (Fig. 2a). In contrast, ~ 25% of establishments from North America into Europe had occurred by 1950, with only a slight indication of convexity in the rate of per-unit-import establishments (Fig. 2b).

**Figure 2.**
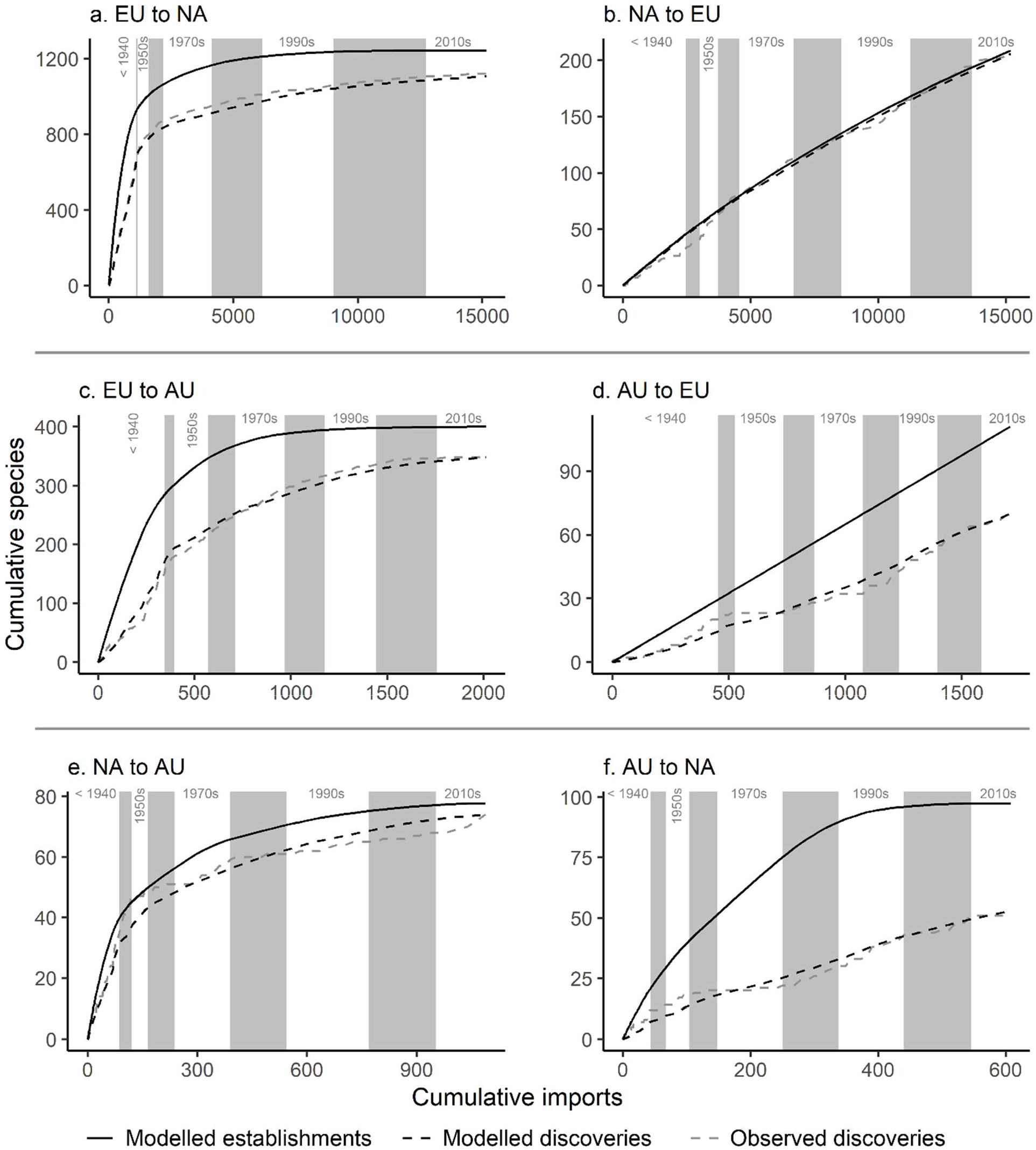
Cumulative discoveries (observed and modelled) and establishments (modelled) of non-native insects exchanged between Europe (EU), North America (NA), and Australasia (AU) versus cumulative import value (inflation-corrected to 2020 British pounds sterling), 1827–2014. Alternating background shading indicates decade boundaries, with shading omitted prior to the 1940s for clarity.

The patterns of cumulative establishments and discoveries between Europe and Australasia resembled those between Europe and North America (compare Figs. 2c, d with Figs. 2a, b). Cumulative plots of establishments and detections of European insects in Australasia are strongly convex, with the best model including a saturation effect. As with the model of European insects into North America, approximately 75% of insect establishments from Europe into Australasia had occurred by 1950 (Fig. 2c). Establishments and discoveries of Australasian insects into Europe showed no effects of saturation, with both observations and models of non-native species exhibiting a linear relationship to imports (Fig. 2d).

Establishments and discoveries of non-native insects between North America and Australasia were within the same order of magnitude in both directions (Fig. 2e, f). There was evidence of saturation in the exchange of non-native insects in both directions between North American and Australasia, though less so from Australasia to North America.

Much like the simple comparisons of total cumulative non-native insect discoveries, the Poisson process models showed strong asymmetries in the exchange of non-native insects between Europe and North America, and between Europe and Australasia.

## Discussion

Considerably more insects have invaded North America and Australasia from Europe than in the opposite directions (Fig. 1). This concurs with the previously observed overrepresentation of tree-feeding insects from Europe in North America^5^, and with non-native insects from the western Palearctic (i.e., Europe) being overrepresented in New Zealand^45^. Our results demonstrate that these asymmetries are consistent across all insect taxa, including non-phytophagous insects.

Asymmetries in establishments between different routes may arise from differences in the size of donor species pools, and thus the numbers of potential invaders^8^. The numbers of described native insects differed significantly in size between Europe, North America, and Australasia (Table 1), but these differences were small compared to the differences in cumulative establishments across the various routes between the three regions. We also caution that small differences in described species richness may be a consequence of differential scientific effort, and not necessarily a true reflection of the ecological community. When we expressed the numbers of non-native insect discoveries exchanged over each route as proportions to the richness of native insects in the donor regions, the resulting coefficient of variation was not noticeably different from that of unadjusted insect discoveries (Fig. 1a, b). We are confident in rejecting the hypothesis of differences in source species pool sizes as a major factor driving the asymmetrical exchange of non-native insects between Europe, North America, and Australasia.

After a non-native species establishes, there is typically a time lag until it is discovered ^46^. Differential discovery lags may lead to asymmetries in cumulative discoveries (but not establishments) between regions. Our models attempted to account for this by estimating the lag between establishment and discovery, allowing us to compare predicted establishment curves among invasion routes. According to these models, the maximum rates of establishment per unit of import value (*r*) of European insects into North America and Australasia were 81 and 19 times greater, respectively, than the establishment rates into Europe from these regions (Table 2). Europe and North America have similar climates and floral composition^47^ and a longer period of colonial contact than between Europe and Australasia^48^, and these factors likely facilitated the relatively greater exchange of insects between Europe and North America.

**Table 2.**
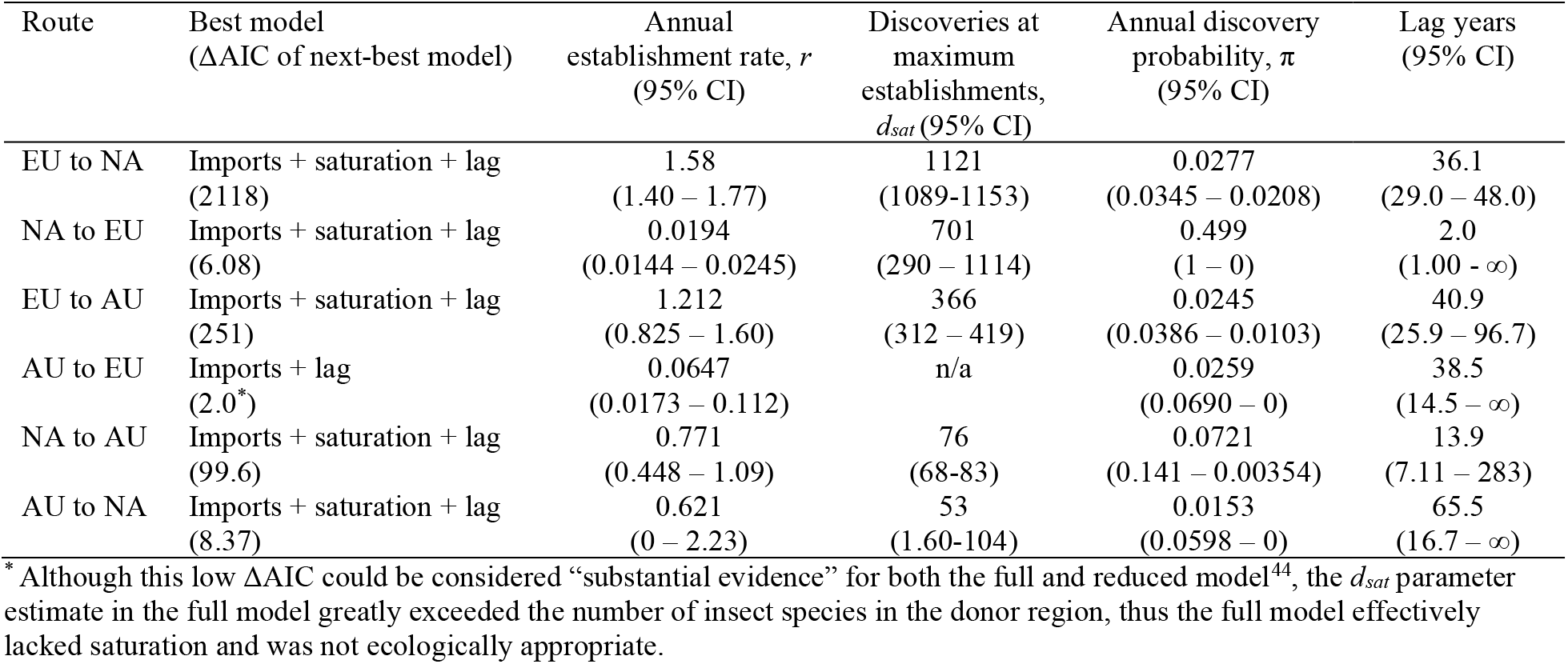
Parameters and 95% confidence intervals of Poisson-process models of establishments and lagged discoveries of non-native insect species exchanged between Europe (EU), North America (NA), and Australasia (AU). All models included a parameter for imports (*r*, the number of annual establishments per billion pounds sterling) and lag (*π*, the annual probability of discovery of established species). Models including an additional term for saturation (a decrease in establishment probability as the cumulative number of discoveries approaches *d_sat_*) were selected for most invasion routes, with model selection based on Akaike information criterion (AIC) values.

The rates of non-native insect establishments between our focal regions have shifted markedly over time. While the overall rate of global establishments of non-native species has not slowed^9,14^, our results show that establishments of European insects in North America and Australasia per unit of import value have drastically decreased since 1950 (Figs. 2a, c). Similarly, Levine and D’Antonio^12^ noted a decline in the rate of accumulation of exotic species per unit of imports in the United States and attributed this to a reduction in the per-ship probability of introducing a species due to the exhaustion of local source species pools. MacLachlan et al.^14^ estimated rates of non-native Hemiptera establishments in the USA over time and found that the risk of species establishment per unit of imports declined from 1850 to 2000, attributing this either to species pool depletion or improved biosecurity effectiveness. Forecasted accumulations of Asian and European bark beetles (Scolytinae) in the USA suggested that the rate of establishments per value of imports will slow over time^10^. There were also declines in the rates of non-native insects exchanged between North America and Australasia, though to a lesser extent (Fig. 2e, f). For North American species established in Australasia, this saturation again starts in the 1950s. For Australasian species established in North America, the evidence for saturation is less convincing, as it is not evident in cumulative discoveries and may be an artifact of an overestimated discovery lag. Discoveries of North American and Australasian species in Europe do not exhibit similarly declining trends and models show only weak or no effects of saturation (Figs. 2b, d). As a result of these trends, the exchange of non-native species per unit of import value between Europe, North America, and Australasia has equalized considerably in recent decades. Most of the asymmetries in discoveries between our focal regions thus accumulated prior to 1950. Although this appears to defy global trends which show accelerating rates of plant and insect invasions after World War II^15^, this is largely a consequence of different ways of measuring establishment rates: as a function of time, or as a function of import value. Declining establishments per unit of import value do not necessarily translate into declining annual establishments, as import values have increased exponentially over time^10^. Should these trends continue, the imbalance of established European insects in North America and Australasia versus the reciprocal exchange will diminish over time. Seebens et al.^49^ also predicted that the rate of non-native arthropod establishments in North America and Australasia would plateau (NA) or slow (AU) through to 2050, while the rate of establishments in Europe would accelerate.

Although we have modelled the declines in per-unit-import establishments as a gradual exhaustion of source species pools, it is also likely that biosecurity measures have contributed. International biosecurity regulations, specifically phytosanitary measures, began in earnest in the latter half of the 20^th^ century^50^. With plant-feeding insects making up 58% of all non-native insect species in our dataset, strengthened phytosanitary measures applying to pathways including live plants, wood, and crops have almost certainly led to contemporary reductions in per-unit-import rates of establishments. Historically, Europe has had relatively weak phytosanitary measures, while Australia and New Zealand have maintained strict phytosanitary policies for many decades^51^. The lack of obvious saturation in the establishments of North American and Australasia insects into Europe may be partly due to Europe’s historically weaker biosecurity. However, differential biosecurity is unlikely to have played a major role in creating the considerable asymmetries we observed in non-native insect establishments and discoveries prior to 1950.

International trade is considered the single most important pathway for unintentional introductions of insects^11^, and greater trade activity generally results in greater propagule pressure of non-native species. Existing literature identifies a positive correlation between import value and the establishment of non-native species^9,12–14^. Similarly, our models provided excellent fits of historical inflation-corrected import values to observed non-native insect discoveries (after accounting for gradual depletion of source pools). However, the modelled establishment rates (r), which represent the maximum rates of establishments per unit of imports prior to any depletion of source pools, differ significantly between the Europe to North America route and its converse, and between the Europe to Australasia route and its converse (Table 2). Furthermore, cumulative import values are nearly identical in both directions between Europe and North America, and only slightly greater from Europe to Australasia than in the opposite direction (Fig. 2a-d). Thus, contrary to our prediction, and despite the important role of trade in facilitating the establishment of non-native species, differences in import values between region pairs cannot explain the observed asymmetries in cumulative discoveries of non-native insects between Europe, North America, and Australasia. This does not mean we can entirely rule out differences in propagule pressure as a driver of asymmetries; import value is a simplistic proxy that does not tell us about the actual nature of the goods exchanged.

Unable to explain the invasional asymmetries between our focal routes as arising from differences in source species pool sizes or trade values, we will now consider how the ecologies of the recipient regions could impact establishment success. Differences in ‘invasibility’ between regions may be driven by differences in the niche diversity available to invaders; plant diversity in the recipient community does, for example, appear to be a strong driver of insect invasion^18^. Niemelä and Mattson^5^ proposed that a history of extensive glaciations may have reduced niche diversity in Europe by leading to extinctions of plant genera. At the time of their publication, Niemelä and Mattson noted approximately 18,000 species of vascular plant in North America (north of Mexico) and 12,000 in Europe, suggesting that this made Europe less invasible to insects^5^. However, Europe has been fairly heavily colonized by insect invaders from regions other than North America, particularly the Asian Palearctic^20^, suggesting that Europe is not strongly resistant to invasion. Additionally, more recent estimates of plant richness show very similar estimates between Europe and North America (Table 1). We find it unlikely that relatively small differences in native host plant richness could lead to such unequal niche opportunities for arriving insects, as observed relationships between plant and insect richness often follow linear or nearly linear relationships^52–54^. When we expressed total discoveries of non-native insects as their proportions to the richness of native vascular plants in the recipient regions, the asymmetry between reciprocal routes, as measured by coefficients of variation, increased slightly (Fig. 1c). This is contrary to predictions that regions with greater native plant diversity would facilitate non-native insect establishments.

Europe stands apart from North America and Australasia in having a long imperial history, including the colonization of the latter two regions. This colonization promoted both deliberate and accidental introductions of European plants^55^. Introductions of exotic plants by colonial powers accelerated in the 19^th^ and early 20^th^ centuries and have left a lasting legacy on the global flora^19^. This is noteworthy, because non-native plant diversity may be a stronger driver of insect invasions than native plant diversity^18^. North America and Australasia each have nearly twice as many extra-continental non-native plant species as Europe^35^ (Table 1). Second to temperate Asia, Europe is the dominant source of non-native plants worldwide, and a considerable proportion of naturalized non-native plants in North America (52%) and Australasia (39%) are native to Europe (Table 1). These patterns are consistent with the observed asymmetries in discoveries of non-native insects between North America, Europe, and Australasia. When cumulative discoveries of non-native insects between donor and recipient regions were expressed as proportions to the number of non-native plants exchanged between the same donor and recipient regions, the asymmetries between invasion routes are considerably reduced (Fig. 1a, d).

The import of European plants into its colonies in North America and Australasia would have promoted the establishment of European insects via two distinct means. First, the import of live plants would have increased the propagule pressure of insects associated with those plants. In the USA, for example, live plant imports may have facilitated the establishment of approximately 70% of damaging non-native forest insects^56^. Second, naturalization of introduced European plants would have provided a landscape containing suitable hosts even to highly host-specific European insects. As a large proportion of herbivorous insects exhibit some degree of host specificity^17^, naturalized non-native plants likely play an important role in facilitating establishments of non-native insects. In Australia, for example, nearly 90% of non-native insect pests and pathogens were associated with non-native plants, only half were associated with native plants, and those associated with native plants were more likely to be polyphagous^23^. The establishment of herbivorous insects from a given region may also be facilitated by the naturalization of host plants from a different region, particularly if those host plants are phylogenetically similar to the insects’ native hosts^11^. In the context of our focal regions, Australasia contains a unique native flora, whereas Europe and North America are united into the Holarctic floral kingdom^47^. Thus, the naturalization of Holarctic plants into Australasia, regardless of their specific origin, may have facilitated both European and North American insect establishments.

## Conclusion

We have documented strong asymmetries in the rates of discoveries of non-native insects between Europe, North America, and Australasia. These asymmetries were strongest prior to 1950 and favored Europe as the dominant source of non-native insects between our focal regions. Our results have allowed us to largely rule out differences in source species pool sizes, overall trade volume (using trade value as a proxy), and native plant diversity as causes of these asymmetries. Although we cannot rule out the possible role of regional differences in insect invasiveness, the introduction of exotic plants driven in large part by European colonization is the most compelling explanation for Europe’s dominance as an ‘exporter’ of non-native insects. This represents both a source of propagule pressure of European insects that was not adequately represented by comparisons of trade value, and changes to floral composition which have facilitated subsequent establishments of European insects. We also cannot rule out other factors not addressed here, such as differences in establishment probability driven by biotic resistance (e.g., native predators and competitors), differences in propagule pressure driven by the specific types of trade goods exchanged between regions, or the effect of establishments originating from non-native populations (‘bridgeheads’). Regardless, we believe our results to be an important step forward in understanding the factors that drive international patterns of non-native species establishments.

## Acknowledgements

This work was supported by the National Socio-Environmental Synthesis Center under funding received from the National Science Foundation DBI-1639145. SBH and DSP were supported by the Natural Sciences and Engineering Research Council of Canada (NSERC) Discovery Grants; RT was supported by USDA Forest Service International Programs 21-IG-11132762-241; AML acknowledges grant EVA4.0, No. CZ.02.1.01/0.0/0.0/16_019/0000803 financed by Czech Operational Programme Science, Research, and Education; AR and EGB were partially supported by the European Union project HOMED (HOlistic Management of Emerging Forest Pests and Diseases-grant No. 771271); CB was supported by Fondation Sandoz-Monique de Meuron and the Swiss National Science Foundation (SNSF); and DR was partially supported by the University of Padua under the 2019 STARS Grants program (project: MOPI—Microorganisms as hidden players in insect invasions).

## Author information

AML, AB, PK, BØ, DR, and DSP developed the initial concept. AML, SBH, and DSP supervised the project. Data collection and database development was performed by AML, RMT, RB, HN, AR, and DSP. Statistical analyses were performed by RLI and TY. The initial draft was prepared by RLI. All authors were involved in the writing and editing process following the initial draft.

## Data Availability

The datasets generated during and/or analysed during the current study are available from the corresponding author on reasonable request.

## References

1. Kenis, M. et al. Ecological effects of invasive alien insects. Biol. Invasions 11, 21–45 (2009).

2. Bradshaw, C. J. A. et al. Massive yet grossly underestimated global costs of invasive insects. Nat. Commun. 7, 12986 (2016).

3. Nahrung, H. F., Liebhold, A. M., Brockerhoff, E. G. & Rassati, D. Forest insect biosecurity: Processes, patterns, predictions, pitfalls. Annu. Rev. Entomol. 68, annurev-ento-120220- 010854 (2023).

4. Blackburn, T. M. et al. A proposed unified framework for biological invasions. Trends Ecol. Evol. 26, 333–339 (2011).

5. Niemelä, P. & Mattson, W. J. Invasion of North American forests by European phytophagous insects. BioScience 46, 741–753 (1996).

6. Visser, V. et al. Much more give than take: South Africa as a major donor but infrequent recipient of invasive non-native grasses. Glob. Ecol. Biogeogr. 25, 679–692 (2016).

7. Vermeij, G. J. An agenda for invasion biology. Biol. Conserv. 78, 3–9 (1996).

8. Vermeij, G. J. Anatomy of an invasion: the trans-Arctic interchange. Paleobiology 17, 281–307 (1991).

9. Seebens, H. et al. No saturation in the accumulation of alien species worldwide. Nat. Commun. 8, 14435 (2017).

10. Liebhold, A. M., Brockerhoff, E. G. & Kimberley, M. Depletion of heterogeneous source species pools predicts future invasion rates. J. Appl. Ecol. 54, 1968–1977 (2017).

11. Brockerhoff, E. G. & Liebhold, A. M. Ecology of forest insect invasions. Biol. Invasions 19, 3141–3159 (2017).

12. Levine, J. M. & D’Antonio, C. M. Forecasting biological invasions with increasing international trade. Conserv. Biol. 17, 322–326 (2003).

13. Lantschner, M. V., Corley, J. C. & Liebhold, A. M. Drivers of global Scolytinae invasion patterns. Ecol. Appl. 30, e02103 (2020).

14. MacLachlan, M. J., Liebhold, A. M., Yamanaka, T. & Springborn, M. R. Hidden patterns of insect establishment risk revealed from two centuries of alien species discoveries. Sci. Adv. 7, eabj1012 (2021).

15. Bonnamour, A., Gippet, J. M. W. & Bertelsmeier, C. Insect and plant invasions follow two waves of globalisation. Ecol. Lett. 24, 2418–2426 (2021).

16. Bertelsmeier, C., Ollier, S., Liebhold, A. & Keller, L. Recent human history governs global ant invasion dynamics. Nat. Ecol. Evol. 1, 1–8 (2017).

17. Forister, M. L. et al. The global distribution of diet breadth in insect herbivores. Proc. Natl. Acad. Sci. 112, 442–447 (2015).

18. Liebhold, A. M. et al. Plant diversity drives global patterns of insect invasions. Sci. Rep. 8, 12095 (2018).

19. Lenzner, B. et al. Naturalized alien floras still carry the legacy of European colonialism. Nat. Ecol. Evol. 6, 1723–1732 (2022).

20. Roques, A. et al. Are invasive patterns of non-native insects related to woody plants differing between Europe and China? Front. For. Glob. Change 2, 91 (2020).

21. Costello, C. J. & Solow, A. R. On the pattern of discovery of introduced species. Proc. Natl. Acad. Sci. 100, 3321–3323 (2003).

22. Turner, R., Blake, R. & Liebhold, A. M. International non-native insect establishment data (0.1). Zenodo (2021) doi:https://doi.org/10.5281/zenodo.5245302.

23. Nahrung, H. F. & Carnegie, A. J. Non-native forest insects and pathogens in Australia: Establishment, spread, and impact. Front. For. Glob. Change 3, 37 (2020).

24. Liebhold, A. M. et al. Invasion disharmony in the global biogeography of native and non-native beetle species. Divers. Distrib. 27, (2021).

25. Mally, R. et al. Moths and butterflies on alien shores: Global biogeography of non-native Lepidoptera. J. Biogeogr. 49, 1455–1468 (2022).

26. GBIF. https://www.gbif.org/ (2022).

27. Blake, R. E. & Turner, R. reblake/insectcleanr. (2021).

28. Fouquin, M. & Hugot, J. Two centuries of bilateral trade and gravity data: 1827-2014. CEPII Work. Pap. 2016–14, (2016).

29. Office for National Statistics. Consumer price inflation time series. https://www.ons.gov.uk/economy/inflationandpriceindices/datasets/consumerpriceindices.

30. McGlone, M. s., Duncan, R. P. & Heenan, P. B. Endemism, species selection and the origin and distribution of the vascular plant flora of New Zealand. J. Biogeogr. 28, 199–216 (2001).

31. Costello, C., Springborn, M., McAusland, C. & Solow, A. Unintended biological invasions: Does risk vary by trading partner? J. Environ. Econ. Manag. 54, 262–276 (2007).

32. Morimoto, N., Kiritani, K., Yamamura, K. & Yamanaka, T. Finding indications of lag time, saturation and trading inflow in the emergence record of exotic agricultural insect pests in Japan. Appl. Entomol. Zool. 54, 437–450 (2019).

33. R Core Team. R: A language and environment for statistical computing. (R Foundation for Statistical Computing, 2021).

34. Byrd, R. H., Lu, P., Nocedal, J. & Zhu, C. A limited memory algorithm for bound constrained optimization. SIAM J. Sci. Comput. 16, 1190–1208 (1995).

35. van Kleunen, M. et al. Global exchange and accumulation of non-native plants. Nature 525, 100–103 (2015).

36. Ulloa, C. U. et al. An integrated assessment of the vascular plant species of the Americas. Science (2017) doi:10.1126/science.aao0398.

37. Arnett, R. H. American insects: A handbook of the insects of America north of Mexico, second edition. (CRC Press, 2000). doi:10.1201/9781482273892.

38. Euro+Med Plantbase. https://www.europlusmed.org/ (2021).

39. de Jong, Y. et al. Fauna Europaea – all European animal species on the web. Biodivers. Data J. e4034 (2014) doi:10.3897/BDJ.2.e4034.

40. Chapman, A. D. Numbers of living species in Australia and the world. (2009).

41. de Lange, P. J. et al. Conservation status of New Zealand indigenous vascular plants, 2017. 87 (2017).

42. Atlas of living Australia. https://www.ala.org.au/ (2022).

43. New Zealand organisms register. https://www.nzor.org.nz/ (2022).

44. Burnham, K. P. & Anderson, D. R. Multimodel inference: Understanding AIC and BIC in model selection. Sociol. Methods Res. 33, 261–304 (2004).

45. Edney-Browne, E., Brockerhoff, E. G. & Ward, D. Establishment patterns of non-native insects in New Zealand. Biol. Invasions 20, 1657–1669 (2018).

46. Essl, F. et al. Socioeconomic legacy yields an invasion debt. Proc. Natl. Acad. Sci. 108, 203–207 (2011).

47. Cox, B. The biogeographic regions reconsidered. J. Biogeogr. 28, 511–523 (2001).

48. Engerman, S. L. & Sokoloff, K. L. Five hundred years of European colonization: Inequality and paths of development. in Settler Economies in World History (eds. Lloyd, C., Metzer, J. & Sutch, R.) 65–103 (Brill, 2013). doi:10.1163/9789004232655.

49. Seebens, H. et al. Projecting the continental accumulation of alien species through to 2050. Glob. Change Biol. 27, 970–982 (2021).

50. Allen, E., Noseworthy, M. & Ormsby, M. Phytosanitary measures to reduce the movement of forest pests with the international trade of wood products. Biol. Invasions 19, 3365–3376 (2017).

51. Eschen, R. et al. International variation in phytosanitary legislation and regulations governing importation of plants for planting. Environ. Sci. Policy 51, 228–237 (2015).

52. Procheş, Ş. et al. Dissecting the plant–insect diversity relationship in the Cape. Mol. Phylogenet. Evol. 51, 94–99 (2009).

53. Basset, Y. et al. Arthropod diversity in a tropical forest. Science (2012) doi:10.1126/science.1226727.

54. Zhang, K. et al. Plant diversity accurately predicts insect diversity in two tropical landscapes. Mol. Ecol. 25, 4407–4419 (2016).

55. Lenzner, B., Essl, F. & Seebens, H. The changing role of Europe in past and future alien species displacement. in From biocultural homogenization to biocultural conservation (eds. Rozzi, R. et al.) 125–135 (Springer International Publishing, 2018). doi:10.1007/978-3-319-99513-7_8.

56. Liebhold, A. M., Brockerhoff, E. G., Garrett, L. J., Parke, J. L. & Britton, K. O. Live plant imports: the major pathway for forest insect and pathogen invasions of the US. Front. Ecol. Environ. 10, 135–143 (2012).

